# Paraventricular Thalamus Neurons Encode Early-life Stress and Execute Consequent Sex-specific Disruptions of Adult Reward Behaviors

**DOI:** 10.1101/2024.04.28.591547

**Authors:** Cassandra L. Kooiker, Mason Hardy, Matthew T. Birnie, Yuncai Chen, Gregory de Carvalho, Amalia Floriou-Servou, Qinxin Ding, Neeraj Thiagarajan, Madison Tetzlaff, Asia Smith, Isabelle T. Yoon, Yahir A. Aranda, Lulu Y. Chen, Tallie Z. Baram

## Abstract

While links between early-life adversity (ELA) and mental illnesses characterized by dysregulated reward behaviors are well-established, the underlying mechanisms remain unclear. In mice, ELA reduces hedonic consumption and interest in sex reward in adult males and, in contrast, augments reward consumption in females. Here, using genetic tagging (TRAPing) we found robust, sex-specific activation of thalamic paraventricular nucleus (PVT) neurons during ELA. Manipulating these neurons in adults normalized reward behaviors: Blocking TRAPed *anterior* PVT neurons restored hedonic consumption in ELA males and augmented hedonic consumption in control females. In contrast, <u>activation</u> of these neurons reduced consumption in control males and ELA females. For posterior PVT, blocking TRAPed cells attenuated excessive reward consumption in ELA females and reduced it in control males. Thus, PVT is key for adaptive brain plasticity; anterior and posterior PVT carry different functions and contribute to the effects of ELA on adult reward behaviors in a sex-dependent manner.

## Introduction

Early-life adversity (ELA) such as abuse, neglect, or societal upheavals impacts the lives of over 50% of the world’s children, with major cost in loss of human potential.^1^ ELA is a strong risk factor for affective disorders including depression, post-traumatic stress disorder (PTSD), and substance use^2–6^, and these risks are modulated by sex. Imaging studies have identified in children and adolescents with a history of ELA the aberrant development of connectivity of reward circuits that are critical for motivation and the experience of pleasure^7–9^.

Complementing human research, animal studies have been pivotal in demonstrating a causal relationship between ELA and disruptions of reward behaviors^10–15^. Such work has also been instrumental in uncovering some of the neurobiological mechanisms involved in the consequences of ELA. For example, experimental animal model studies have shed light on how ELA alters subsequent responses to stress and how a transient ELA is converted into lasting outcomes via persistent changes in gene expression through epigenetic mechanisms^10,16–18^.

The influence of ELA on reward behaviors is modulated by sex, as the relation of ELA to subsequent emotional problems differs in men and women^19–21^. As examples, women with a history of early-life trauma are more likely than men to crave comfort food and develop eating-and opioid use disorders^22–24^. In contrast, men with a history of ELA are more likely to develop alcohol use disorder. Similarly, animal models of ELA, including our widely used resource scarcity paradigm, lead to comparable sex-modulated outcomes^25–31^, providing a platform for uncovering the underlying mechanisms. However, while significant progress has been made, key questions remain unanswered: Where in the brain is ELA encoded? Do neuronal populations engaged by ELA also contribute to the disruption of adult reward behaviors? And what is the neurobiological basis for the profound sex-differences in these dysregulations?

Using activity-dependent genetic tagging we have previously identified robust activation of the paraventricular nucleus of the thalamus (PVT) during the first postnatal week in a model of ELA in the mouse^32,33^. Notably, among all analyzed brain regions, the PVT was the only region that distinguished ELA from standard rearing as quantified by the number and type of cells activated: in males, a larger number of PVT neurons were genetically tagged, and, in both sexes, a higher proportion of tagged cells expressed the receptor for the stress-sensitive peptide corticotropin releasing hormone (CRH) in ELA mice as compared to typically reared mice. These findings suggest the PVT may encode and hence potentially mediate the long-term behavioral consequences of ELA.

The PVT is an important component of limbic and emotional processing networks^34^. It contributes to reward-seeking behaviors by integrating information on internal states as well as the external environment and incorporating prior salient experiences^35–37^. The PVT interconnects with brain regions implicated in stress and aversive experiences, such as extended amygdala, brainstem, and medial prefrontal cortex (mPFC), and projects densely to the nucleus accumbens (NAc), a critical node of the reward circuit^38–41^. Via this unique connectivity, the PVT straddles competing motivational circuits to gate the expression of motivated behaviors and modulate responses to stress^42–47^. However, whereas the role of the PVT in incorporating past experiences into decisions involving motivated reward behaviors has been established^48–50,37^, it has remained unknown whether such past experiences can be as remote as those taking place neonatally. Further, the roles of distinct domains of this heterogeneous structure in integrating prior experiences to influence reward behavior, as well as potential sex-differences in these functions, have remained unexplored. The current studies address these fundamental information gaps.

## Results

### Differential activation of PVT by early-life adversity versus typical neonatal experiences

We capitalized on the TRAP2 mouse system, which allows for permanent activity-dependent labeling of neurons active during a defined 24-36 h time-period (**Fig.1A**)^51^. We imposed ELA on groups of mice using the limited bedding and nesting (LBN) model established in our lab and adopted widely^52,27^. The paradigm involves rearing pups for a week during an early sensitive developmental period (postnatal days [P]2-9) in a resource-scarce environment^14,52^, which consistently induces sex-dependent changes in reward behaviors including hedonic consumption of food later in life^11,13,27^. We activated the genetic tagging (TRAP2) system midway through the ELA epoch (P6), thereby inducing recombination (“TRAPing”) in neurons active on P6 and P7. Quantification of TRAPed PVT neurons showed that ELA led to an augmented number of activated PVT neurons compared to standard rearing in males (**Fig. 1B,C; Extended Data Fig. 1**), in accord with our prior finding^32^. In females, whereas the total number of activated cells was similar in ELA and control mice (**Fig. 1B, D; Extended Data Fig. 1**), a larger proportion of PVT neurons active during ELA expressed the principal receptor for corticotropin releasing hormone, CRFR1, as was also the case in males (**Fig 1E-G**). Because ELA did not increase the total number of CRFR1^+^ neurons within the PVT of either male or female mice (**Fig 1H,I**), this indicated that the augmented proportion of CRFR1^+^ TRAP neurons in the PVT of ELA mice resulted from preferential engagement of the population of CRFR1^+^ neurons among all PVT cell types.

**Figure 1.**
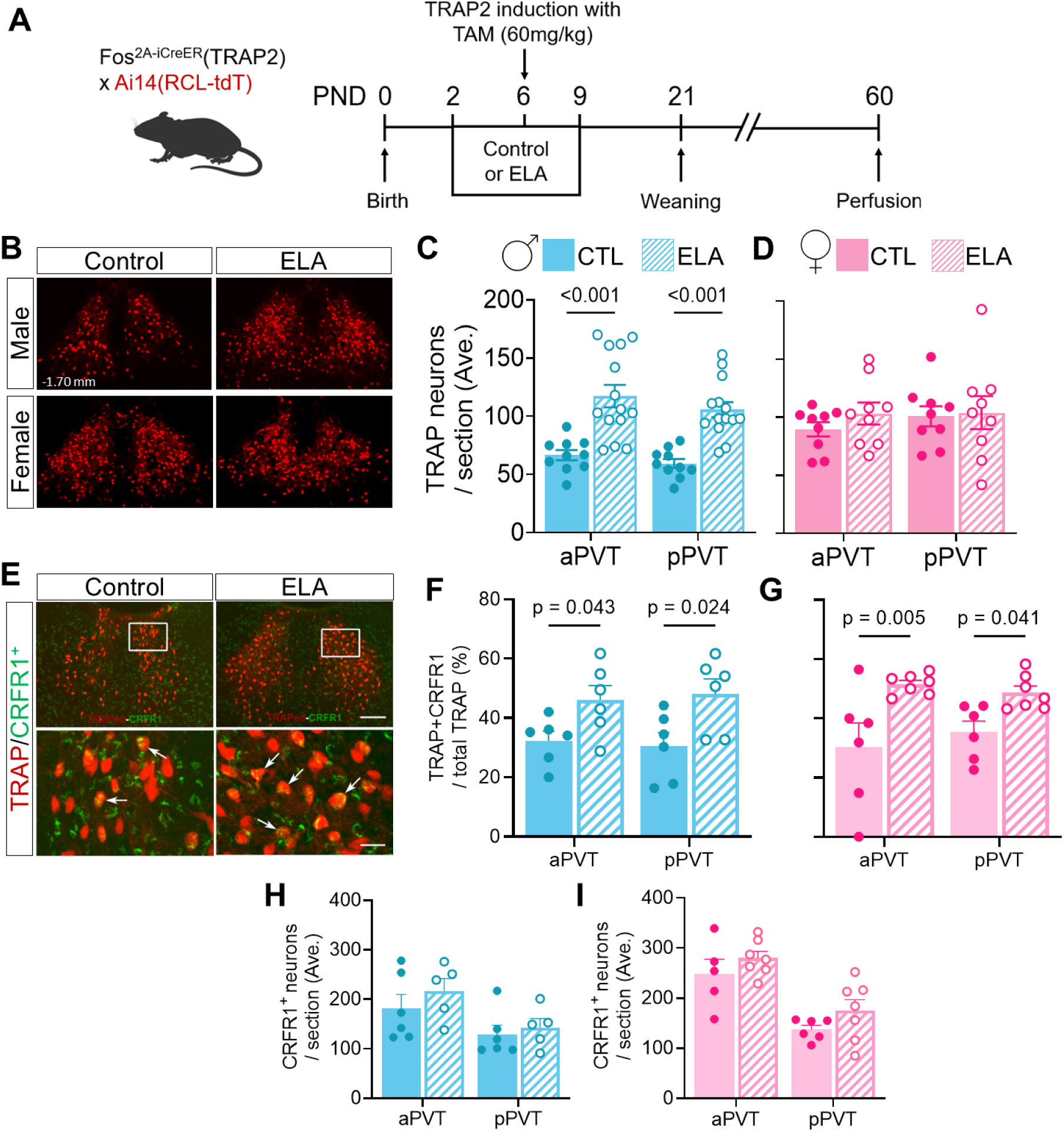
ELA-induced PVT activation is sex-dependent. (**A**) Experimental timeline of TRAP2 studies. (**B**) Representative images from male control (top left), male ELA (top right), female control (bottom left) and female ELA (bottom right) TRAP2 mice showing tdTomato reporter expression in the posterior PVT following TRAP induction with tamoxifen on P6. (**C,D**) Quantification of tdTomato^+^ neurons in male (**C**) and female (**D**) control and ELA mice. ELA augments early-life PVT activation in males (n = 10-14 mice/group; F_1,44_ = 42.92, p <0.001, main effect of rearing, followed by Holm-Šídák’s multiple comparisons *post-hoc* tests: anterior PVT control males vs anterior PVT ELA males p<0.001; posterior PVT control males vs posterior PVT ELA males p<0.001; 2-way ANOVA). No significant difference in early-life PVT activation was identified in females following ELA (n = 9 mice/group; F_1,32_ = 0.686, p = 0.414, main effect of rearing; 2-way ANOVA). (**E**) Representative images from male control (left) and male ELA (right) TRAP2 mice showing tdTomato reporter expression colocalized with CRFR1 expression in the posterior PVT following TRAP induction with tamoxifen on P6. Higher magnification images are shown in the bottom panels. (**F,G**) Quantification of the colocalization of CRFR1 with tdTomato^+^ neurons in a subset of male (**F**) and female (**G**) control and ELA mice following immunofluorescent staining for CRFR1. ELA increased TRAP-CRFR1 colocalization in males (n = 6 mice/group; F_1,20_ = 12.14, p = 0.002, main effect of rearing, followed by Holm-Šídák’s multiple comparisons *post-hoc* tests: anterior PVT control males vs anterior PVT ELA males p = 0.043; posterior PVT control males vs posterior PVT ELA males p = 0.024; 2-way ANOVA) and in females (n = 6-7 mice/group; F_1,22_ = 15.79, p<0.001, main effect of rearing, followed by Holm-Šídák’s multiple comparisons *post-hoc* tests; anterior PVT control females vs anterior PVT ELA females p = 0.005; posterior PVT control males vs posterior PVT ELA males p = 0.041; 2-way ANOVA). (**H,I**) Quantification of CRFR1^+^ PVT neurons in male (**H**) and female (**I**) control and ELA mice. ELA did not increase the number of CRFR1^+^ neurons in males (n = 5-6 mice/group; F_1,18_ = 1.106, p = 0.307, main effect of rearing; 2-way ANOVA) or females (n = 5-7 mice/group; F_1,21_ = 3.200, p = 0.088, main effect of rearing; 2-way ANOVA). Bars represent mean ± SEM.

### Inhibition of anterior PVT neurons active early in life normalizes reward behaviors in adult ELA males

Exposure to ELA led to sex-dependent, opposing disruptions of adult reward behaviors (**Fig. 2A)**. Adult male ELA mice had an anhedonia-like phenotype, whereby they consumed less of a palatable food (i.e., Cocoa Pebbles cereal; high-sugar, low nutrients) than control males (**Fig. 2C**) and were less interested in a sex cue (i.e., the scent of a female in heat)^27,53^ (**Extended Data Fig. 2**). In contrast, adult ELA females exhibited augmented reward consumption relative to females reared in standard cages^54–56^ (**Fig. 2D**). The influence of ELA was reward-specific rather than driven by metabolic needs, as both consumption of chow and body weight were not influenced by rearing (**Extended Data Fig. 3A,B**). Additionally, ELA did not alter anxiety or general locomotor behavior (**Extended Data Fig. 3C,D**). The divergence of the impact of ELA on adult behaviors in male and female mice suggested sex-differences in the processes encoding ELA and governing its influence on adult hedonic behaviors. Because the anterior and posterior PVT differ in cell composition and connectivity, endowing them with both overlapping as well as distinct and even opposing functions^57–60^, we tested the possible separate roles of the anterior and posterior PVT in mediating the consequences of ELA in male and female mice.

**Figure 2.**
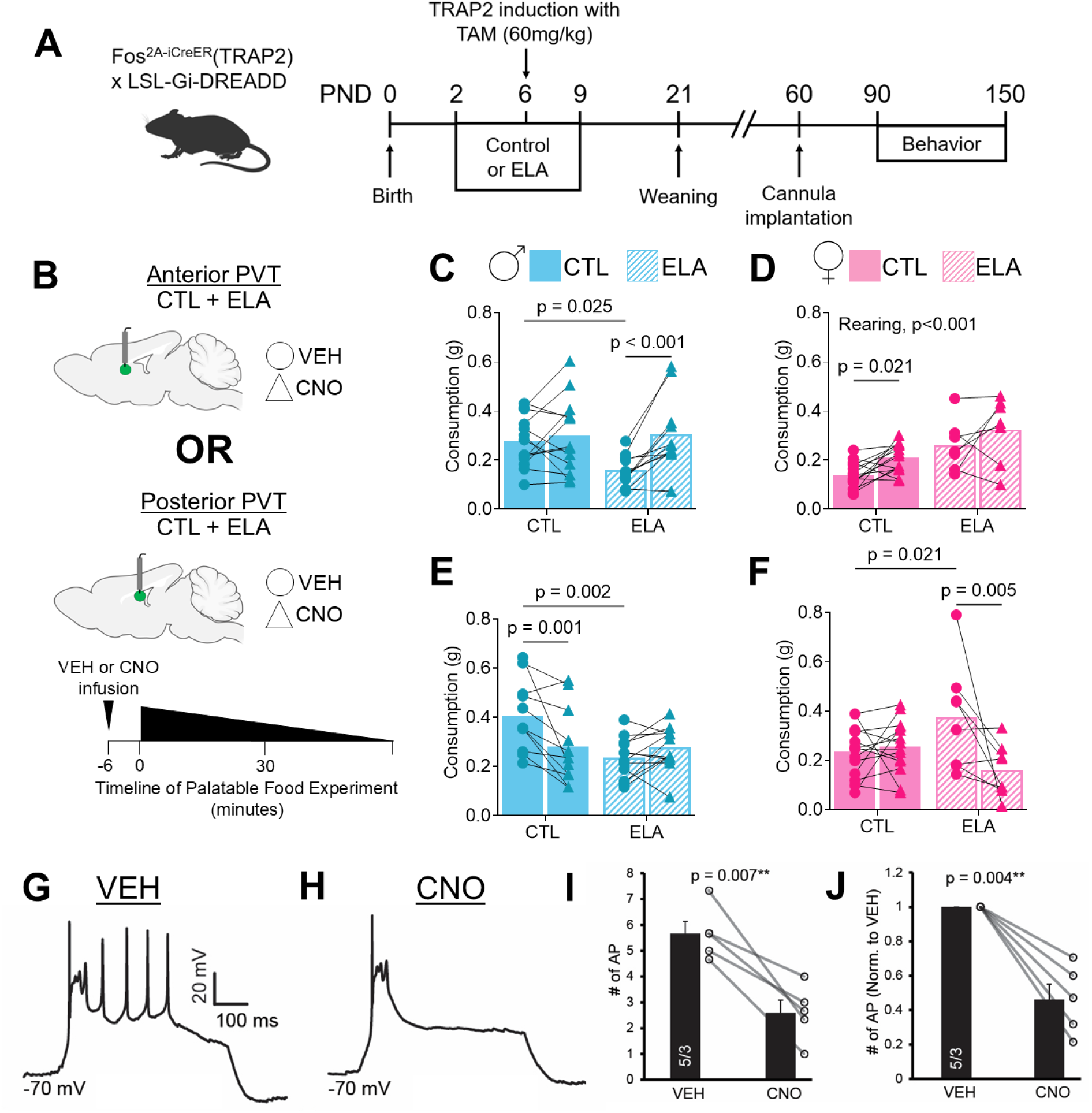
TRAPed anterior PVT neurons are responsible for the anhedonia-like phenotype in ELA males, and TRAPed posterior PVT cells are involved in females. (**A**) Experimental schematic for chemogenetic inhibition of TRAPed PVT neurons. TRAP2 x LSL-Gi-DREADD mice received a tamoxifen injection on P6, inducing expression of an inhibitory DREADD in neurons active during ELA or standard rearing. A cannula for drug infusion was implanted in the aPVT or pPVT of P60 mice and palatable food consumption was measured following vehicle or CNO infusion. (**B**) Experimental schematic for anterior PVT inhibition and timeline of PF test. (**C**) Palatable food consumption following intra-aPVT vehicle or CNO microinfusion in control and ELA males (n = 11-13 mice/group; F_1,22_ = 6.045, p = 0.022, main effect of rearing x treatment; F_1,22_ = 11.62, p = 0.003 main effect of treatment, followed by Fisher’s LSD multiple comparison *post-hoc* test for rearing and treatment: vehicle CTL vs vehicle ELA males p=0.025; vehicle ELA vs. CNO ELA males p<0.001 2-way RM ANOVA). (**D**) Palatable food consumption following intra-aPVT vehicle or CNO microinfusion in control and ELA females (n = 7-14 mice/group; F_1,19_ = 16.09, p<0.001, main effect of rearing; F_1,19_ = 7.651, p = 0.012, main effect of treatment, followed by Fisher’s LSD multiple comparisons *post-hoc* tests of treatment: vehicle CTL vs. CNO CTL females p = 0.021; 2-way RM ANOVA). (**E**) Palatable food consumption following intra-pPVT vehicle or CNO microinfusion in control and ELA males (n = 11-12 mice/group; F_1,21_ = 13.01, p = 0.002, main effect of rearing x treatment, followed by Fisher’s LSD *post-hoc* tests for rearing and treatment: vehicle CTL vs vehicle ELA males p=0.002; vehicle CTL vs. CNO CTL males p=0.001; 2-way RM ANOVA). (**F**) Palatable food consumption following intra-pPVT vehicle or CNO microinfusion in control and ELA females (n = 8-14 mice/group; F_1,20_ = 7.724, p = 0.012, main effect of rearing x treatment; F_1,20_ = 5.137, p = 0.035 main effect of treatment, followed by Fisher’s LSD *post-hoc* tests rearing and treatment: vehicle CTL vs vehicle ELA females p=0.021; vehicle ELA vs. CNO ELA females p=0.005; 2-way RM ANOVA). (**G**) Representative trace of a PVT neurons expressing hM4Di in the vehicle condition recorded in current clamp; slow current was applied to hold the membrane at -70 mV, and a depolarizing current step (500 ms) was applied. Intensity of current step was set so that the neuron evoked 4-6 action potentials. (**H**) Representative trace of action potentials from an a hM4Di-expressing PVT neuron after clozapine-N-oxide (CNO) superfusion showing a decrease in number of action potentials. (**I**) Summary graph showing a significant decrease in total number of action potentials in hM4Di-expressing PVT neurons (pairwise t-test: VEH: 5.67 ± 0.46, CNO: 2.6 ± 0.49; p = 0.007). (**J**) Summary graph showing a significant decrease in number of action potentials in hM4Di-expressing PVT neurons normalized to the vehicle condition (pairwise t-test: VEH: 1 ± 0, CNO: 0.46 ± 0.09; p = 0.004). Bars represent mean ± SEM.

First, we tested if selective chemogenetic inhibition of ELA-activated (TRAPed) anterior PVT neurons influenced the reduced hedonic consumption in adult ELA males(**Fig. 2B)**. We used Fos^2A-iCreER+/-^::R26-LSL-Gi-DREADD mice that were reared in either standard or ELA cages and injected with tamoxifen midway through the ELA or control rearing period (on postnatal day [P]6). Tamoxifen administration to these mice led to persistent expression of the inhibitory designer receptor exclusively activated by designer drugs (DREADD) HA-hM4Di-pta-mCitrine only in neurons active from P6-P7, enabling chemogenetic inhibition of this neuronal population later in life. To limit chemo-inhibition to PVT neurons only, we implanted in adult mice cannulae in either the anterior or posterior PVT and administered the DREADD ligand clozapine-N-oxide (CNO) via the cannulae. Following infusion of CNO or vehicle, male mice underwent one of two reward tasks: palatable food consumption during a one-hour free access period, or exploration of a sex cue (the scent of a female in heat)^61,62^. Inhibition of ELA-engaged neurons in the anterior PVT of adult male ELA mice restored their palatable food consumption back to levels seen in typically reared mice (**Fig. 2C**) and also tended to increase interest in the sex cue (**Extended Data Fig. 2A).** By contrast to ELA males, inhibition of TRAPed anterior PVT neurons had no effect on control males in either task (**Fig. 2C, Extended Data Fig. 2A)**. Testing for potential off-target effects of CNO, we found no changes in locomotion attributable to CNO or rearing (**Extended Data Fig. 4A**).

In females, selective inhibition of TRAPed anterior PVT neurons increased palatable food consumption (**Fig. 2D**), reaching significance only in <u>control</u> females. The absence of a significant effect on consumption in ELA females following this manipulation may reflect a ‘ceiling effect’ in the ability to drive further consumption in this group. These data suggest that early-life activated cells in the anterior PVT (or this region as a whole) may act as a brake on hedonic food consumption. Again, CNO did not influence locomotion in either female group (**Extended Data Fig. 4B**).

Together, although these studies have not excluded potential inhibition of axon terminals from afferent projections to PVT of TRAPed neurons in other regions, the findings demonstrate the contribution of anterior PVT neurons activated during ELA to the resulting anhedonia-like behaviors in males and suggest that ELA influences anterior PVT neurons differentially by sex.

### Inhibition of TRAPed posterior PVT neurons normalizes hedonic consumption in adult ELA females

The posterior PVT is activated by adverse experiences in adults and influences behaviors in response to subsequent stress^18,43–45,48^. Therefore, we determined if inhibition of posterior PVT neurons activated by stress early in life (ELA) would ameliorate the consequent disruption of adult reward behaviors.

In adult ELA males, selective chemogenetic inhibition of TRAPed posterior PVT neurons had no effect on their diminished consumption of palatable food (**Fig. 2E**), nor did it alter the attenuated interest in the sex cue preference task (**Extended Data Fig. 2C**). In contrast, in control males, inhibiting TRAPed posterior PVT neurons decreased palatable food consumption (**Fig. 2E**), consistent with a ‘pro-hedonic role for these cells. The effect of selective chemogenetic inhibition of posterior PVT neurons activated early in life was highly sex-specific. In adult control females, inhibition of these neurons had no influence on palatable food consumption. Whereas in ELA females, CNO infusion into the posterior PVT significantly reduced their aberrantly augmented palatable food consumption, reversing the effect of ELA (**Fig. 2F**) and again, consistent with a pro-hedonic function of these cells. Following these experiments, we confirmed that chemogenetic inhibition of PVT neurons with hM4Di indeed suppressed neuronal firing. Using whole cell patch clamp recordings (**Fig. 2G-J**), application of CNO robustly inhibited Gi-expressing neurons, measured by a significant reduction in evoked action potentials (**Fig. 2I,J**). These results support the contribution of ELA-engaged neurons in the posterior PVT to the aberrantly augmented reward consumption consequent to ELA in females and, together with the results in control males, suggest that posterior PVT cell activity may promote hedonic consumption.

### Distinct, sex-specific and rearing-dependent functions of anterior and posterior PVT neurons

The results of the above experiments selectively inhibiting anterior or posterior PVT neurons in males and females reared typically or in ELA were surprising. They suggested both sex- and rearing-specific roles of early-life-engaged neurons within these PVT subregions that conform to the operational model shown in **Fig. 3G**. The model posits that TRAPed <u>anterior</u> PVT neurons act as a brake on reward behaviors, and their augmented function in ELA males contributes to reduced hedonic consumption. In contrast, TRAPed <u>posterior</u> PVT neurons facilitate reward behaviors, and their augmented early-life activation in ELA females contributes to the augmented hedonic consumption in this group. Accordingly, inhibiting these neurons in adulthood reduces palatable food consumption in control males, but not in the already ‘anhedonic’ ELA males. In addition, ELA influences these two PVT sub-regions differentially by sex. In males, ELA enhances the function (or “weight”) of anterior PVT neurons, contributing to diminished adult reward behaviors. In contrast, persistently augmented function of TRAPed posterior PVT neurons drives aberrantly increased hedonic consumption in adult ELA females.

**Figure 3.**
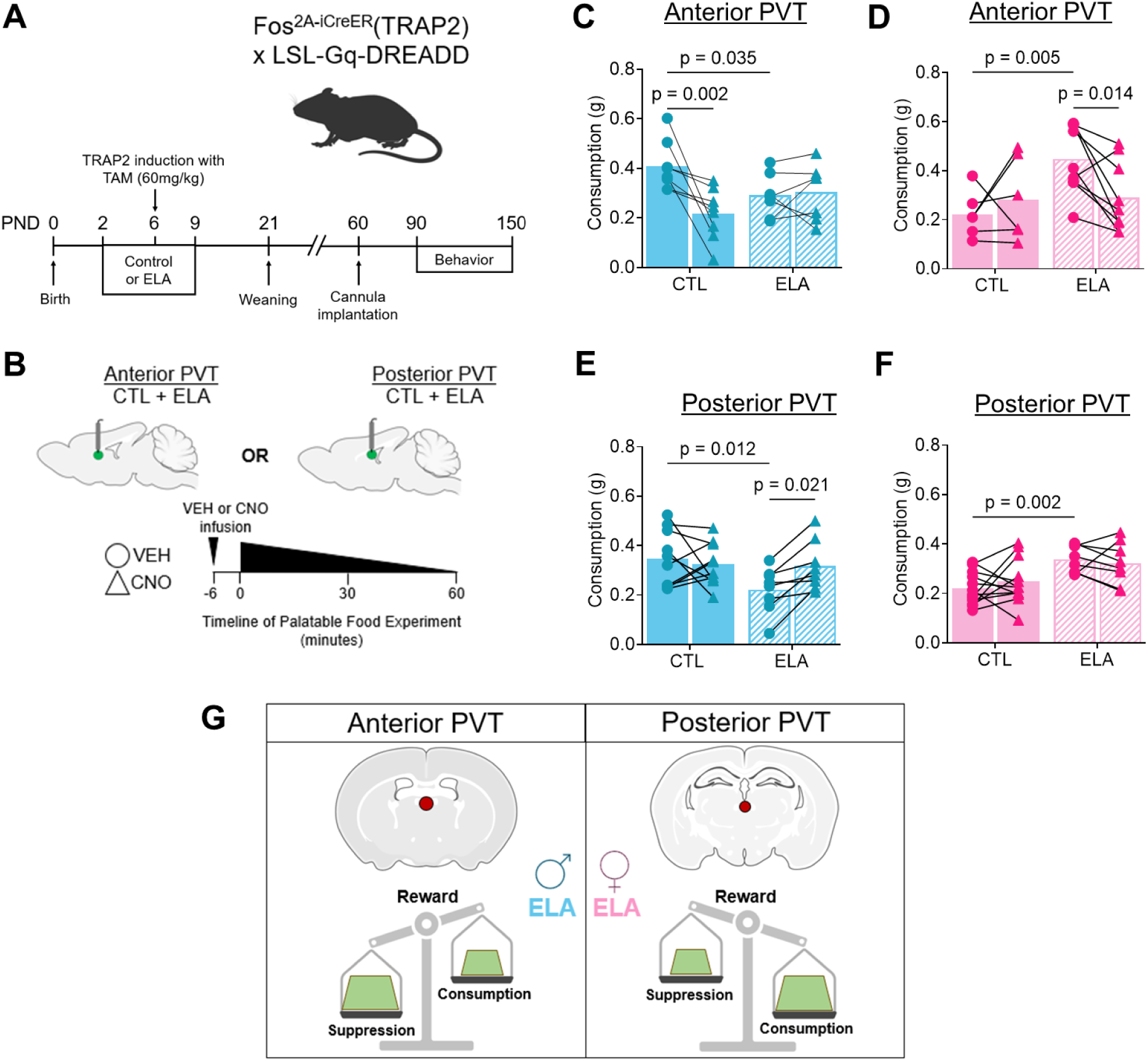
Excitation of TRAPed anterior PVT neurons induces anhedonia-like behavior, whereas activation in posterior PVT neurons promotes reward. (**A**) Experimental schematic for chemogenetic excitation of TRAPed PVT neurons. TRAP2 x CAG-LSL-Gq-DREADD mice received a tamoxifen injection on P6, inducing expression of an excitatory DREADD in neurons active during ELA or standard rearing. A cannula for drug infusion was implanted in the aPVT or pPVT of P60 mice and palatable food consumption was measured following vehicle or CNO infusion. (**B**) Experimental schematic for anterior PVT excitation and timeline of PF test. (**C**) Palatable food consumption following intra-aPVT vehicle or CNO microinfusion in control and ELA males (n=7-8 mice/group; F_1,13_ = 8.339, p=0.013, main effect of rearing x treatment; F_1,13_ = 6.280, p=0.026, main effect of treatment, followed by Fisher’s LSD multiple comparison *post-hoc* test for rearing and treatment: vehicle CTL vs vehicle ELA males p = 0.035; vehicle CTL vs CNO CTL males p=0.002; 2-way RM ANOVA). (**D**) Palatable food consumption following intra-aPVT vehicle or CNO microinfusion in control and ELA females (n = 6-9 mice/group; F_1,13_ = 6.155, p=0.028, main effect of rearing x treatment, followed by Fisher’s LSD *post-hoc* tests for rearing and treatment: vehicle CTL vs vehicle ELA females p=0.005; vehicle ELA vs CNO ELA females p=0.014; 2-way RM ANOVA). (**E**) Palatable food consumption following intra-pPVT vehicle or CNO microinfusion in control and ELA males (n = 8-10 mice/group; F_1,16_ = 5.508, p=0.032, main effect of rearing x treatment, followed by Fisher’s LSD *post-hoc* tests for rearing and treatment: vehicle CTL vs vehicle ELA males p=0.012; vehicle ELA vs CNO ELA males p=0.021; 2-way RM ANOVA). (**F**) Palatable food consumption following intra-pPVT vehicle or CNO microinfusion in control and ELA females (n = 8-12 mice/group; F_1,18_ = 12.24, p=0.003, main effect of rearing, followed by Fisher’s LSD multiple comparison *post-hoc* test for rearing: vehicle CTL vs vehicle ELA females p=0.002; 2-way RM ANOVA). (**G**) Hypothesized roles of anterior and posterior PVT neurons TRAPed in early-life, and the impact of rearing and sex. In control mice, females have greater activity (“weight”) of the anterior PVT, which suppresses reward, whereas males have augmented activity of the posterior PVT, which promotes reward. Following exposure to ELA, the “weight” of anterior and posterior PVT influence appears to invert between sexes, such that ELA males have augmented anterior PVT activation and ELA females exhibit greater posterior PVT activation.

The proposed model generates testable hypotheses. Specifically, it predicts that *<u>activating</u>* anterior PVT neurons in control males and ELA females will repress reward behaviors, whereas activating posterior PVT neurons will, instead, promote reward behaviors in ELA males and control females. We performed experiments to test these hypotheses in Fos^2A-iCreER+/-^::CAG-LSL-Gq-DREADD mice (**Fig. 3A,B**) and found that, as expected, chemogenetic activation of TRAPed <u>anterior</u> PVT neurons reduced both palatable food consumption and the interest in female urine in control male mice, but not in ELA mice (**Fig. 3C, Extended Data Fig. 2C**). In females this excitation reduced hedonic consumption in the ELA group, in accord with the model’s prediction (**Fig. 3D**). Activation of the posterior PVT augmented palatable food consumption and interest in sex cues in ELA males as expected (**Fig. 3E, Extended Data Fig. 2D**). However, posterior PVT TRAP excitation did not increase palatable food consumption in control females (**Fig. 3F**), indicating that the role of early-life activated posterior PVT cells in hedonic consumption may be confined to the ELA group. Alternatively, posterior PVT cells were not sufficiently activated by the control rearing environment to lead to enduring impact on reward behaviors.

### Mechanisms of the enduring effects of ELA on adult reward behaviors may involve a reactivation of PVT neurons active early in life

The experiments described above show that inhibiting selectively PVT neurons activated early in life is sufficient to reverse the effect of ELA on male and female mice in a region-specific manner. These findings suggest that the same neurons TRAPed during the ELA period are responsible for executing the effects of ELA on reward behaviors. This scenario predicts that PVT neurons active during ELA should be **re**activated during adult reward behaviors, measurable by cFos expression. We tested for the reactivation of these neurons by exposing TRAP2::Ai14 ELA and control mice (which had received tamoxifen on P6 to permanently visualize neurons activated early in life) to 1.5 hours of free access to palatable food, a time period optimal for visualizing cfos expression with immunohistochemistry^84^ (**Fig 4A**). Following perfusion and brain processing, we quantified cFos expression and the co-localization of cFos with the TRAP reporter, indicating a reactivation of neurons engaged early in life (**Fig 4B**).

**Figure 4.**
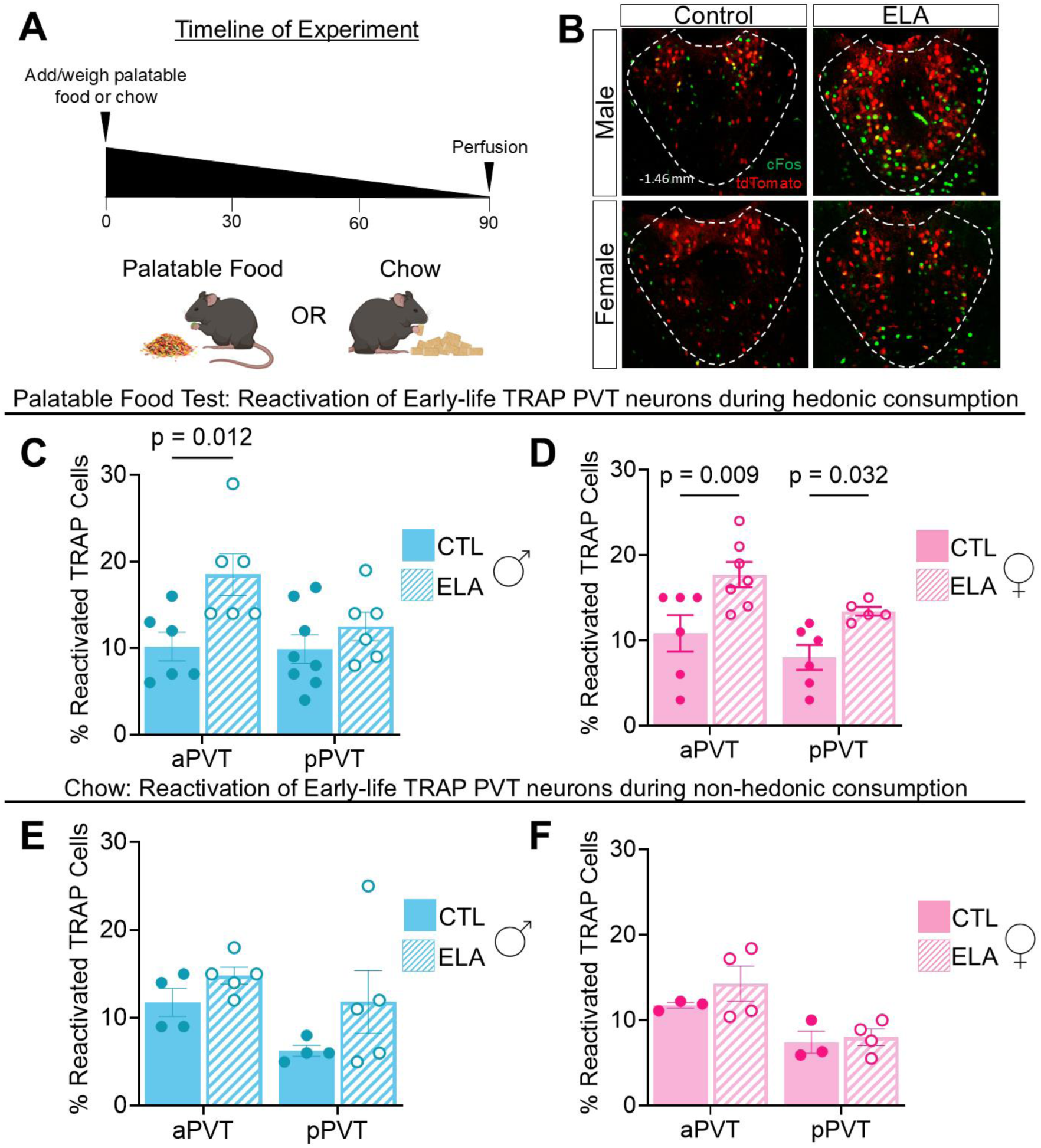
Reward-induced reactivation of early-life-engaged PVT neurons. (**A**) Schematic of the experimental timeline for palatable food and chow TRAP reactivation. (**B**) Representative images of cFos expression and tdTomato (i.e., TRAPed neurons) co-expression in the PVT of control (left) and ELA (right) mice immediately following 1.5 hrs. of free access to palatable food. (**C**) Percent reactivation of early-life TRAPed PVT neurons in control and ELA males following 1.5 hrs. of free access to palatable food (n = 6-8 mice/group; F_1,22_ = 8.544, p = 0.008, main effect of rearing, followed by Holm-Šídák’s multiple comparisons test: anterior PVT control vs anterior PVT ELA, p=0.012; 2-way ANOVA). (**D**) Percent reactivation of early-life TRAPed PVT neurons in control and ELA females (n = 5-7 mice/group; F_1,20_ = 5.080, p=0.036, main effect of region; F_1,20_ = 15.00, p<0.001, main effect of rearing, followed by Holm-Šídák’s multiple comparisons test for rearing: anterior PVT control vs anterior PVT ELA, p=0.009; posterior PVT control vs posterior PVT ELA, p=0.032). (**E**) Percent reactivation of early-life TRAPed PVT neurons in control and ELA males following 1.5 hrs. of free-access consumption of regular chow (n = 4-5 mice/group; F_1,14_ = 3.636, p=0.077, main effect of rearing; F_1,14_ = 3.722, p=0.074, main effect of region; 2-way ANOVA). (**F**) Percent reactivation of early-life TRAPed PVT neurons in control and ELA females following 1.5 hrs. of free-access consumption of regular chow (n = 3-4 mice/group; F_1,10_ = 1.163, p=0.306, main effect of rearing; F_1,10_ = 13.26, p=0.005, main effect of region; 2-way ANOVA). Bars represent mean ± SEM.

In males, reactivation of *anterior* but not *posterior* PVT cells distinguished ELA and control mice (**Fig. 4B,C**), as expected. In females, reactivation was more robust in both PVT regions of adult ELA mice (**Fig. 4B,D**). ELA-induced augmentation of neuronal reactivation was specific to the consumption of rewarding food, as there was no difference between ELA and control mice in response to regular chow (**Fig 4E,F**). In addition, reactivation during reward exposure of neurons in the lateral hypothalamic area and the ventromedial hypothalamus, both regions associated with feeding, did not differ significantly among the groups (**extended data Fig. 5**). Together, these data support the notion that exposure to ELA leads to sex- and rearing-specific activation of neurons in both the anterior and posterior PVT, and to augmented reactivation of the same neurons by adult reward consumption in the ELA adults.

## Discussion

While our prior work identified the PVT as a brain region that distinguishes early-life stress from typical experiences^32^, the functional implications of the region’s selective activation have remained unknown. The current studies used chemogenetic manipulations and single-cell resolution measures of activity to show that, in addition to encoding ELA, the PVT executes the dysregulated adult reward behaviors resulting from ELA in a manner that depends upon sex, rearing, and PVT region. In general, anterior PVT neurons genetically tagged during ELA mediate anhedonia-like reward behavior deficits in ELA males, while, in contrast, TRAPed posterior PVT neurons enable aberrantly augmented hedonic consumption in females. Using chemogenetic excitation, we demonstrate that activation of anterior PVT neurons can induce an ELA-like phenotype (i.e., anhedonia) in control males as well as attenuate the augmented reward consumption in ELA females. Additionally, excitation of the posterior PVT can normalize reward consumption in ELA males in a manner similar to the effects of anterior PVT inhibition. Thus, anterior PVT neurons activated during early-life seem to act as brakes on reward behaviors while posterior PVT neurons activated during early-life tend to enhance them, such that the sex-specificity of the effects of ELA stems from differential ELA recruitment or weighting of anterior PVT in males and posterior PVT in females. Together, these discoveries position the PVT as a major hub of early-life plasticity and as a potential target for addressing the maladaptive plasticity induced by ELA.

### The PVT both encodes early life experiences and contributes to their consequences

The use of genetic tagging allowed qualitative and quantitative testing of which brain regions express Fos in response to distinct environmental stimuli during the first week of life in the mouse. In addition, the methods enabled addressing whether, in adult mice, the same neurons are necessary and sufficient to generate dysregulated reward behaviors observed in adult ELA mice.

We first reaffirmed that, among all brain regions analyzed, the PVT is unique in its differential activation by ELA versus typical rearing conditions. Notably, the hippocampal formation, the canonical memory-encoding brain region, was minimally and equally active in control and ELA mice. This observation is consistent with the developmental timing of ELA, occurring prior to the debut of operational hippocampus-mediated memory systems. This takes place during the third and fourth postnatal weeks^59,60^, leaving unexplained how the brain encodes experiences during a window of time in early life prior to the formation of semantic memories. Given that experiences in the early postnatal period can have profound impacts on emotional health in adulthood, a different brain region or regions may be responsible for encoding these experiences. Our studies point to the PVT as an important participant in this process.

Next, we chemogenetically manipulated the same PVT cells active early in life, and established that blocking them, in a sex- and PVT region-specific manner was sufficient to reverse the dysregulated reward behaviors in adult ELA mice. Further, a sex-specific contribution of anterior vs posterior PVT was apparent. Accordingly, selective <u>activation</u> of anterior PVT TRAPed neurons during adult reward behaviors, repressed hedonic consumption in control males, though the expected augmentation of hedonic consumption in control females upon activation of posterior PVT TRAPed neurons was not observed. That is, in females posterior PVT TRAPed neurons are necessary for augmented hedonic consumption following ELA, but not sufficient to increase hedonic consumption in typically reared females. Together, these findings indicate that the same cells activated during the first week of life by the distinct rearing environments are responsible for the aberrant adult reward behaviors. Notably, while selective manipulation of the TRAPed cells was sufficient to modulate reward behaviors, our data do not exclude the possibility that all cells, TRAPed or not, may participate in attenuating or promoting these behaviors in a sex and region dependent manner.

### Distinct roles for anterior and posterior PVT in the sex-specific ELA-induced disruptions of reward behaviors provide insights into the functions of these PVT regions

In adult mice, we tested whether the population of PVT neurons activated by ELA contributes to behavioral disruptions following ELA. Specifically, ELA led to significant reductions in palatable food consumption in males and the augmentation of palatable food consumption in females. While we observed modest variability in palatable food intake between cohorts, likely due to subtle strain differences between lines (i.e., TRAP2 crossed with Gi- or Gq-expressing lines) and/or different experimenters, the overall effect of ELA was consistent (and has been found in numerous mouse strains as well as in rats)^11,13,27,28,31, 63^.

Inhibition of TRAPed neurons in the <u>anterior</u> PVT of adult ELA males ameliorated their reduced consumption of palatable food, whereas excitation recapitulated the ELA phenotype in control males, suggesting that these neurons may drive the anhedonia-like disruption of reward behaviors consequent to ELA in males. In contrast, inhibition of anterior PVT TRAPed neurons in ELA females had no effect on their augmented hedonic consumption. These data support a role for anterior PVT neurons as ‘brakes’ on hedonic consumption and increased functional weight of these neurons in ELA males.

In females, by contrast, inhibition of TRAPed posterior PVT neurons ameliorated ELA-induced augmented consumption of palatable food whereas inhibition of the same neuronal populations in ELA males had no effect. Additionally, posterior PVT inhibition reduced reward consumption in control males, while posterior PVT excitation rescued anhedonia-like behavior in ELA males. These data support a ‘pro-hedonic’ role for the posterior PVT neurons activated early in life in both males and females, and suggest that their aberrant function may contribute to the dysregulated hedonic consumption in ELA females.

The discoveries emerging from probing the role of anterior and posterior PVT in disrupted reward behaviors induced by ELA provide insights into general functional principles of the anterior and posterior PVT. For the anterior PVT, inhibition of the TRAPed neurons augmented palatable food consumption in control females, consistent with the elimination of a ‘brake’ on reward behaviors. This was not seen in control males, perhaps reflecting a ‘ceiling effect’. Blocking posterior PVT TRAPed neurons in males reduced reward behaviors in the controls, consistent with elimination of a pro-reward-drive. No effect was observed in control females in whom consumption was already low.

These data suggest that PVT neurons activated early in life (and potentially all neurons) contribute to either promoting (posterior PVT) or attenuating (anterior PVT) reward behaviors, and these contributions are further modulated by sex and rearing. Such distinct regional functional roles of PVT neurons are consistent with the existing body of work in adult rodents: The anterior PVT has roles in arousal, reward suppression, and motivational conflict. Activation of anterior PVT neurons can inhibit reward-seeking, induce aversion, and suppress food consumption^60,74^, whereas inhibition increases reward-seeking and reduces anxiety-like behavior^74,75^. During motivational conflict between reward approach and threat avoidance, inhibition and excitation of CRH-expressing neurons in the anterior PVT biases behavior toward approach or avoidance, respectively^76^. Additionally, neurons in the anterior PVT show diurnal fluctuations in depolarization/excitability^77^ and are modulated by sleep–wake transitions^78^. Cell-type-specific analyses identify *Ntrk1*-positive and *Drd2*-negative neurons as predominant cell types in the anterior PVT, with *Drd2*-negative cells being modulated by arousal and reward approach, and excitation of *Ntrk1* cells leading to reduced reward consumption^58,60,79^. In contrast, *Drd2*-positive neurons predominate in the posterior PVT, and neuronal activity in this region has been linked to the encoding of aversive stimuli, expression of avoidance behavior, the formation of both threat-and reward-cue associations, and signaling reward approach and acquisition^47,49,58,80,81^. Excitation of posterior PVT neurons increases food and sucrose consumption^60,75,82^, and plasticity of PFC inputs to the posterior PVT supports the maintenance of reward-seeking behavior^47^.

A potential basis for the opposing behavioral roles of anterior and posterior PVT involves their distinct patterns of efferent connectivity to cortical and limbic regions (for a comprehensive review, see Kirouac, 2025). Although both PVT regions innervate the infralimbic cortex (IL), prelimbic cortex (PL), ventral subiculum (vSub), nucleus accumbens (NAc), amygdala, and bed nucleus of the stria terminalis (BNST), they do so with markedly different density patterns^39,41,83^. The anterior PVT more densely innervates the IL, PL, dorsomedial NAc shell (dmNAcSh), and vSub, whereas the posterior PVT preferentially targets the ventromedial NAc shell (vmNAcSh), NAc core (NAcC), central amygdala (CeA), basolateral amygdala (BLA), and lateral BNST^38,39,41,83^. Recent work suggests distinct cell-type-specific efferents of the PVT^60,76^, lending additional complexity and nuance to the complex interplay among PVT regions, cell-type gradations, and efferent connectivity patterns along the anteroposterior axis that collectively shape behavioral responses to rewarding and aversive stimuli. The anterior and posterior PVT appear to exert both coordinated and opposing influences on appetitive and aversive behavior, and disruption of the balance between these subregions may contribute to dysregulated behavior. While these processes do not account for the observed sex differences, we recently identified, using brain-wide mapping, sex-dependent innervation patterns of cell-type-specific projections^63^. Such variation in synaptic connectivity might also exist in the PVT, contributing to the sex-modulated effects of ELA on reward behaviors.

### Potential mechanisms transducing transient ELA into enduring changes in adult behavior

Our genetic tagging captured cells expressing Fos, which is both a marker of neuronal activation as well as an immediate early gene that robustly influences transcriptional programs^84–87^. Thus, exposure to ELA leads to activation of neurons in the PVT followed by persistent changes in these neurons via transcriptional mechanisms. Indeed, recent work from our group shows sex-dependent changes in gene expression at baseline and a strikingly sex-dependent reward-induced activation of gene translation in ELA mice as compared to controls^71^. These lasting changes in ELA-engaged PVT neurons may alter their functions during motivated behaviors, as supported by the chemogenetic studies presented here.

Support for an enduring effect of ELA on TRAPed cells is apparent from the studies assessing their reactivation during adult reward consumption. Reactivation of genetically tagged neurons has also been shown in the NAc, following a similar ELA paradigm^88^. Here, more TRAPed cells were reactivated in ELA vs control mice, consistent with increased responsiveness to reward consumption. This may arise from synaptic alterations^89–91^, epigenomic/transcriptional changes^84–87^, or both. Additionally, ELA may cause a shift in the composition of cell populations activated during the early postnatal period, such that the ensembles of neurons TRAPed in ELA mice include cells that would otherwise not be engaged in controls. This may result from patterns of sensory input not typically present in control conditions, resulting from the observed aberrant maternal care-derived sensory signals characteristic of this paradigm^14,92^. A potential circuit mechanism for the differential activation of PVT neurons in control and ELA mice is the afferent input into these neurons. Indeed, we have recently found that stress-sensitive CRH-expressing neurons projecting from the Barrington nucleus (BN) reach the PVT and are poised to activate CRFR1^+^ neurons^93^. The disrupted patterns of maternal care behaviors in ELA cages^14^ include fragmented bouts of licking and grooming of the anogenital region. These signals may be conveyed through the BN, a region that elicits sex-dependent behaviors^94,95^. Finally, ELA may induce alterations in the developmental trajectory of the PVT, which could similarly result in a unique composition of early-life activated neuronal populations. Such shifts in the maturation of specific cell types following the disruption of early-life sensory input have been observed within the hypothalamus^96^, providing a plausible mechanism through which early-life experiences bias the engagement of specific cells. Notably, evidence for the differing composition of TRAPed cells comes from the augmented CRFR1^+^ TRAPing in ELA males and females. In sum, altered cell types or synaptic connectivity of the TRAPed neurons may render them more responsive to reward, as measured here by Fos expression during an adult reward task.

In conclusion, the present studies position the PVT as a major hub of early-life plasticity. They show a role for PVT neurons active during ELA in mediating enduring disruption of reward behaviors, which are sex dependent. We identify new distinct and opposing roles for anterior and posterior PVT and pinpoint the anterior PVT as a primary contributor to disrupted reward behaviors in adult males and the posterior PVT in females, suggesting intrinsic sex-dependent differences in the operations of the reward circuit that are operational already in the neonatal period. Thus, the PVT constitutes a key potential target for addressing the maladaptive plasticity induced by ELA.

## Materials and Methods

### Mice

Fos^tm2.1(icre/ERT2)Luo^/J (Fos^2A-iCreER^; Jax #030323), B6.129-Gt(ROSA)^26Sortm1(CAG-CHRM4*,-mCitrine)Ute^/J (R26-LSL-Gi-DREADD; Jax #026219), B6N;129-Tg(CAG-CHRM3*,-mCitrine)1Ute/J (CAG-LSL-Gq-DREADD; Jax #026220), and B6.Cg-Gt(ROSA)26Sortm14(CAG-tdTomato)Hze/J (Ai14; Jax #007914) mice were obtained from The Jackson Laboratory or bred in house. All mice were housed in a temperature-controlled, quiet, and uncrowded facility on a 12-hour light/dark schedule (lights on at 7:00 AM, lights off at 7:00 PM) prior to cannula surgery. After surgery, mice were maintained on a 12-hour reverse light cycle (lights on at midnight, lights off at noon) to enable behavioral testing during the dark-phase (active period); we have previously found no difference in any behavioral test between normally housed and reverse light cycle–housed mice when tested at the same point in their circadian cycle. After cannula implantation, mice were single-housed and handled frequently to mitigate potential behavioral changes related to single-housing stress. We observed no appreciable behavioral differences between group-housed or single-housed ELA (or control) mice. Mice were provided with ad libitum access to water and a global soy protein-free extruded diet (2020X Teklad; Envigo). Progeny of Fos^2A-iCreER^ mice bred with Ai14 reporter mice were used for immunohistochemistry studies. Progeny of Fos^2A-iCreER^ mice bred with R26-LSL-Gi-DREADD or CAG-LSL-Gq-DREADD mice were used for behavioral studies. All experiments were performed in accordance with National Institutes of Health guidelines and were approved by the University of California Irvine Institutional Animal Care and Use Committee .

### Limited Bedding and Nesting (LBN) Model of Early Life Adversity (ELA)

Mouse dams were checked every morning for presence of copulatory plugs while paired with a male and placed into single housing on embryonic day (E)17, after which point the cage was checked for presence of pups every 12 hours. On the morning of postnatal day (P)2, litters were culled to a maximum of 8 (minimum of 4) including both sexes, and the ELA paradigm was initiated as previously described^97,98^. Control dams and pups were placed in cages with a standard amount of corn cob bedding and one 5 x 5 cm cotton nestlet. ELA dams and pups were placed in a cage atop a fine-gauge aluminum mesh (cat #4700313244; McNichols Co., Tampa, FL) set approximately 2.5 cm above the cage floor, which was covered with a thin layer of corn cob bedding. ELA dams received one-half nestlet. All cages were placed in rooms with robust ventilation to avoid accumulation of ammonia. Both ELA and CTL dams and pups were left undisturbed until the afternoon of P6 at which point the pups were briefly removed from the home cage, placed in a new cage on a heating pad, and injected subcutaneously with 60 mg/kg tamoxifen (catalog no. T5648; MilliporeSigma) dissolved in corn oil (catalog no. C8267; MilliporeSigma) as previously described^32^. Pups were then returned to their home cage and left undisturbed until the morning of P10. On the morning of P10, each pup was weighed, and the dam and pups were moved to standard cages. P6 was chosen as the time point for tamoxifen injections because it is roughly at the midpoint of the ELA paradigm, allowing for optimal tagging of neurons activated by ELA rearing.

### Cannula implantation surgery

Adult mice (P60+) were anesthetized with 1-3% isoflurane and placed in a stereotactic frame. For intra-PVT drug infusions, a guide cannula (23G, 30G obturator) was implanted above the aPVT (cannula placement at AP: -0.4 mm; ML: -0.5 mm; DV: 2.6 mm, 8° angle toward midline; terminal point of injector at AP: -0.4mm, ML: 0.0 mm, DV: 3.6 mm) or pPVT (cannula placement at AP: - 1.6 mm; ML: -0.5 mm; DV: 1.9 mm, 8° angle toward midline; terminal point of injector at AP: -1.6 mm, ML: 0.0 mm, DV 2.9 mm) and mounted to the skull with a screw and dental cement. Mice were given at least two weeks to recover before beginning behavioral experiments.

### Brain slice preparation for whole cell patch clamp recordings

Mice (P120 - 140) were deeply anesthetized with isoflurane and quickly decapitated. Acute coronal slices (300 μm) were obtained using a vibratome (Leica V1200S, Deer Park, IL) in ice-cold cutting solution containing (in mM): 228 Sucrose, 11 glucose, 26 NaHCO3, 1 NaH2PO4, 2.5 KCl, 5 sodium ascorbate, 3 sodium pyruvate, 10 MgSO4, 0.5 CaCl2 (305-310 mOsm, pH 7.2). Slices equilibrated in a homemade chamber for 20 – 30 mins at 31° C then 30 min at room temperature in artificial cerebrospinal fluid (aCSF) containing (in mM): 119 NaCl, 26 NaHCO3, 1 NaH2PO4, 2.5 KCl, 11 glucose, 1.3 MgSO4-7 H2O, and 2.5 CaCl2 (290 - 295 mOsm, pH 7.4) before being transferred to a recording chamber. aCSF in the equilibration chamber additionally contained 5 mM Ascorbic acid and 2 mM sodium pyruvate. aCSF flow rate during recording was set to 3 ml/min for all recordings.

### Whole-cell patch-clamp recordings

Whole-cell patch-clamp recordings were obtained from PVT neurons expressing AAV5-hSyn-hM4D(Gi)-mCherry (Addgene plasmid # 50475; RRID:Addgene_50475). AAV5-hSyn-hM4D(Gi)-mCherry was injected into the anterior PVT (AP: -0.4 mm; ML: -0.5 mm; DV: 3.65 mm, 8° angle toward midline) and pPVT (AP: -1.6 mm; ML: -0.5 mm; DV: 2.9 mm, 8° angle toward midline) and expressed for ∼6 weeks before recording. Cells were visualized under IR-DIC using an upright BX51WI microscope (Olympus, Breinigsville, PA). Data were collected with a Multiclamp 700B amplifier, Digidata 1550B, and pClamp11 (Molecular Devices, San Jose, CA). All recordings were acquired in voltage clamp (unless stated otherwise) at 34°C and were low pass filtered at 2 kHz and digitized at 10 kHz. Recording pipettes were filled with internal solution containing (in mM): 130 K-Gluconate, 4 NaCl, 10 HEPES, 0.25 EGTA, 10 phosphocreatine, 4 MgATP, 0.3 NaGTP (295 – 305 mOsm, pH 7.4 with KOH). The calculated liquid junction potential was 14.8 mV, and this was not corrected during recordings. All pipettes (3 - 4 MΩ) were pulled from borosilicate glass (Narishige PC-100, Amityville, NY). Access resistance (Ra) was monitored at the beginning and end of the recording and cells with an increase of > 20% in Ra were discarded. For all recordings, a constant current was applied to the neuron to hold the membrane potential at -70 mV. After obtaining whole cell configuration, depolarizing steps (500 ms) applied through the recording pipette were used to evoke action potentials. The size of the step (pA) was set to the amount of current that evoked 4 – 6 action potentials. If the neuron was unable to evoke 4 action potentials it was discarded and not considered for analysis. Three sweeps of depolarizing current step were recorded, then 1 µM clozapine-N-oxide (CNO) was applied through superfusion. Three depolarizing sweeps were recorded again after CNO adequately washed in the recording chamber. Analysis consisted of comparing the number of action potentials evoked before and after superfusion of CNO. We also normalized the number of action potentials after CNO superfusion to the number of action potentials in the vehicle condition for comparison of relative change of the number of action potentials. All data was compared using a pairwise t-test with an α-critical of 0.05.

### Chemogenetic studies

For excitatory (TRAP2 x CAG-LSL-Gq-DREADD mice) and inhibitory (TRAP2 x R26-LSL-Gi-DREADD) DREADD experiments, clozapine-N-oxide (CNO) or vehicle (artificial cerebrospinal fluid, aCSF) were administered via microinfusion through an intra-PVT cannula. CNO (cat #HB6149, HelloBio, UK) was dissolved in aCSF at a concentration of 1 mM, and 0.2 µl of solution was administered at a rate of 50 nl/min. Microinjection needles were left in place for 2 min. following injection to allow for diffusion.

### Behavioral assays

Behavioral assays were carried out in low light (<15 lux) and during the animal’s active period. Prior to the start of testing, all mice were placed in the behavior room for at least 1 hour to acclimate to their surroundings. For all chemogenetic experiments (**Fig. 2 and Fig. 3**), studies were carried out using within-subject design. All mice were tested twice for each test (palatable food and sex cue), receiving vehicle or CNO infusion through guide cannulae. Vehicle and CNO were counterbalanced across the first and second test days for each experiment. A minimum of 48 hours between testing was included for CNO washout. Sex cue and palatable food test order were counterbalanced. These studies included 8 cohorts (anterior or posterior PVT; excitatory or inhibitory TRAP-DREADD; male or female) tested separately. Each cohort consisted of age-, genotype-, and sex-matched control and ELA mice (i.e., 16 groups total).

### Palatable food and chow consumption

Single-housed mice were habituated to ∼1 g Cocoa Pebbles cereal in the home cage for 1 hour on each of three days preceding the test day. On the test day, intra-PVT CNO or vehicle was administered 8 minutes prior to placing ∼1 g pre-weighed Cocoa Pebbles into the cage. After leaving mice undisturbed for 1 hour (chemogenetic studies) or 1.5 hours (TRAP reactivation study), uneaten Cocoa Pebbles were collected and weighed to determine consumption. Mice that never consumed >100 mg across all habituation and test days were excluded. For tests of TRAP reactivation, 1.5 hours was selected to optimize cFos expression: peak cFos protein expression occurs 1-2 hours following neuronal activation^100^. In mice assigned to the chow group, consumption of chow was measured by weighing chow 1.5 hours prior to perfusion, and again at the time of perfusion.

### Sex cue preference

For experiments in males, urine was collected from female mice in the estrous phase of their estrous cycle. Urine was collected on the same day as the test and was stored in capped 0.2 ml PCR tubes at 4°C until use. 60 µl of urine and 60 µl of almond extract (a control, non-rewarding scent; Pure Almond extract, Target) were pipetted onto separate cotton swabs and affixed within opposite corners of the home cage of single-housed males. Intra-PVT CNO or vehicle were administered immediately before the test. During the test, mice were given free access to both swabs for 3 min. Time spent with the nose within 3 cm of each swab and locomotion were tracked using 3-point rodent tracking in Ethovision XT15 software (Noldus, US). Data was analyzed both for effects of CNO on swab interaction (sex cue vs. almond odor) and locomotion (total distance traveled).

### Open Field Test: Anxiety & Locomotion

Following habituation to the behavior room for 60+ minutes, TRAP mice were tested in an open field for baseline levels of locomotion and anxiety-like behavior. Time spent in the center and periphery of the open field arena, as well as total distance traveled (i.e., locomotion) were recorded using 3-point rodent tracking in Ethovision XT15 software (Noldus, US).

### Brain processing and analysis

Following behavioral studies, mice were euthanized with Euthasol and transcardially perfused with ice cold phosphate-buffered saline (PBS) (pH = 7.4) followed by 4% paraformaldehyde in 0.1M sodium phosphate buffer (pH = 7.4). Perfused brains were postfixed in 4% paraformaldehyde in 0.1M PBS (pH= 7.4) for 12-16 hours before cryoprotection in 15%, followed by 30% sucrose in PBS. Brains were then frozen and coronally sectioned at a thickness of 25 µm (1:5 series) using a LeicaCM1900 cryostat (Leica Microsystems). For the purposes of cell quantification, the anterior PVT is defined as -0.34mm to -0.94mm posterior to Bregma, and the posterior PVT at -1.46mm to -2.18mm. Tissue from cannulated mice was assessed for location of cannula placement, and mice were excluded if the terminal point of the cannula was determined to be significantly outside of these pre-defined regions (see **Extended Data Fig. 6** for injector terminal points). For quantification of TRAP cells, CRFR1^+^ cells, and Fos, sections were excluded from analysis if tissue was significantly damaged during the collection process or if quality of tissue impeded accurate imaging and quantification.

### Immunocytochemistry

For fluorescent immunolabeling, free-floating brain tissue sections were washed and permeabilized in 0.01 M PBS containing 0.3% Triton X-100 (PBS-T) for 15 minutes (3 x 5 minutes). Sections were then incubated in 5% normal donkey serum (NDS) for 1 hour. Next, sections were incubated at 4°C for 72 hours in relevant primary antibodies: Goat anti-CRFR1 (1:2,000) or Rabbit anti-cFos (Oncogene, Ab-5, PC38). Secondary antibody labeling was then performed at room temperature, in the dark, with agitation using either 1:400 Donkey anti-Goat Alexa Fluor 488 (Invitrogen, CAT# A-11055) or 1:400 Goat anti-Rabbit Alexa Fluor 488 (CAT #A-11008, Invitrogen, US) for 2 hours at room temperature. Tissue was washed 3 x 5 minutes in PBS-T, mounted on gelatin-coated slides, and coverslipped with Vectashield mounting medium with DAPI (Vectashield, Cat # H-1200, Vector Laboratories).

Goat anti-CRFR1 (Everest Biotech, CAT# EB08035; RRID:AB_2260976) is a polyclonal antibody that binds to the N-terminus of CRFR1 (amino acids 107-117), a sequence that is distinct from that of the CRFR2 ^101,102^. Immunolabeling of free-floating mouse brain tissue using Goat anti-CRFR1 strongly recapitulates established cellular characteristics and distribution of CRFR1^+^ neurons throughout the brain^103–106^. Additionally, this antibody has been validated in Western blots of human, mouse, and rat tissue lysates, where it produces immunoblots of the appropriate molecular weight^106^ (manufacturer’s datasheet), matching previously reported results from the use of a separate CRFR1 antibody targeting the C-terminus^103^.

### Image acquisition

Confocal images were collected using an LSM-510 confocal microscope (Zeiss) with an apochromatic 10X, 20X, or 63X objective. Virtual z sections of 1 µm were taken at 0.2- to 0.5-µm intervals. Image frame was digitized at 12-bit using a 1024 X 1024 pixel frame size.

### Statistical analysis

CRFR1, tdT^+^ and cFos^+^ neuron numbers were counted manually in Fiji^99^. All quantifications and analyses were performed using stereological principles including systematic unbiased sampling and without knowledge of group assignment. Statistical analyses were carried out using GraphPad Prism (GraphPad Software). To examine significance of cell number differences throughout the anteroposterior axis of the PVT, we used two-way ANOVA with Holm-Šidák’s *post-hoc* tests. For chemogenetic experiments, all analyses were two-way repeated measures (RM) ANOVAs comparing the effects of treatment (vehicle or CNO) and rearing (CTL or ELA) with Fisher’s LSD *post-hoc* tests. Due to the opposing effects of ELA on male and female reward behavior, groups were analyzed within sex.

## Author contributions

C.L.K., M.H. M.T.B., and T.Z.B. designed the study. C.L.K., M.H., Q.D., N.T., I.T.Y., and Y.A.A. contributed to data collection. C.L.K. and M.H. conducted all surgeries. C.L.K., M.H., Q.D., N.T., Y.A.A. conducted all behavior studies. C.L.K., M.H. and N.T. conducted histological analyses. C.L.K., M.H., and T.Z.B. wrote the paper. All authors discussed and commented on the manuscript. We thank Graciella Angeles, Amber Aourangzeb, Mackenzie Morrison, and Di Wu for technical assistance.

## List of Supplementary materials

a. Statistics Table ANOVA
b. Statistics Table Effect Size
c. Extended Data:

Extended Data Figure 1. Representative images of early-life TRAP throughout the PVT

Extended Data Figure 2. Inhibition and excitation of early-life TRAPed PVT neurons during the sex cue test

Extended Data Figure 3. ELA does not influence anxiety, locomotion, regular chow consumption or adult body weight

Extended Data Figure 4. Intra-PVT CNO infusion and rearing do not influence locomotion

Extended Data Figure 5. TRAP reactivation does not differ between control and ELA mice in the lateral hypothalamus or ventromedial hypothalamus

Extended Data Figure 6. Representative map of PVT injection locations for DREADD experiments

