## supplemental for "Paraventricular Thalamus Neurons Encode Early-life Stress and Execute Consequent Sex-specific Disruptions of Adult Reward Behaviors"

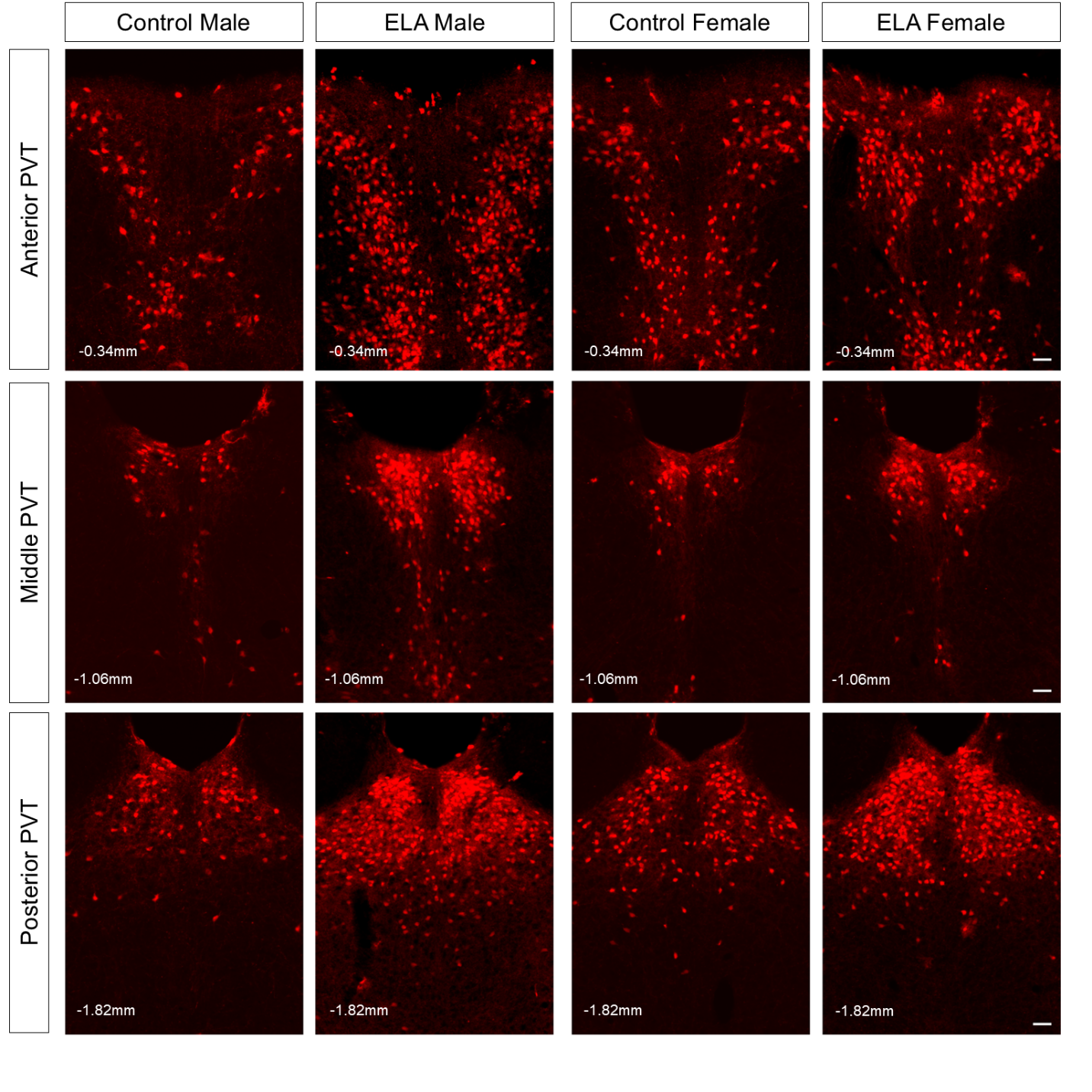

**Extended Data Fig 1. Representative images of early-life TRAP throughout the PVT.** P6/7 TRAP-tdTomato expression in the anterior PVT, middle PVT, and posterior PVT (from top to bottom rows) of CTL male, ELA male, CTL female, and ELA female (left to right). Scale bar = 50 microns.

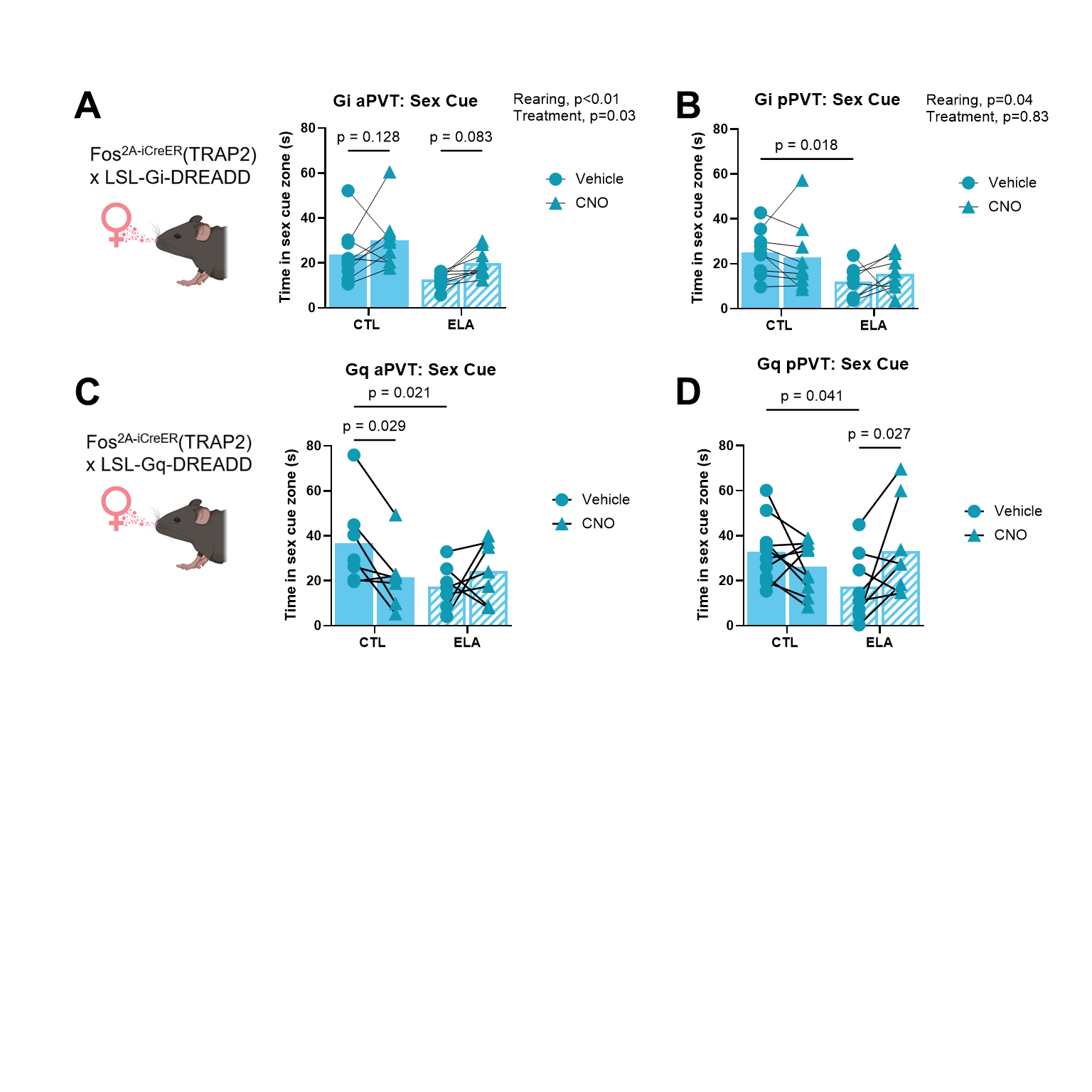

**Extended Data Figure 2. Inhibition and excitation of early-life TRAPed PVT neurons during the sex cue test.** (**A**) Time spent sniffing estrous female urine in a 3 min. test following intra-aPVT vehicle or CNO microinfusion in TRAP2 x LSL-Gi-DREADD control and ELA males. (n = 9-9 mice/group; F_1,16_ = 8.995, p = 0.009, main effect of rearing; F_1,16_ = 5.952, p = 0.027, main effect of treatment, followed by Fisher’s LSD multiple comparison *post-hoc* tests for treatment: vehicle CTL vs CNO CTL p = 0.128; vehicle ELA vs CNO ELA p = 0.083; 2-way RM ANOVA. (**B**) Time spent sniffing estrous female urine in a 3 min. test following intra-pPVT vehicle or CNO microinfusion in TRAP2 x LSL-Gi-DREADD control and ELA males. (n = 8-9 mice/group; F_1,15_ = 4.925 p = 0.042, main effect of rearing, followed by Fisher’s LSD multiple comparison *post-hoc* test: vehicle CTL vs. vehicle ELA p = 0.018; 2-way RM ANOVA). (**C**) Time spent sniffing estrous female urine in a 3 min. test following intra-aPVT vehicle or CNO microinfusion in TRAP2 x LSL-Gq-DREADD control and ELA males (n = 7 mice/group; F_1,12_ = 6.573, p = 0.025, main effect of rearing x treatment, followed by Fisher’s LSD *post-hoc* tests for rearing and treatment: vehicle CTL vs. CNO CTL p = 0.029; vehicle CTL vs vehicle ELA p = 0.021; 2-way RM ANOVA). (**D**) Time spent sniffing estrous female urine in a 3 min. test following intra-pPVT vehicle or CNO microinfusion in TRAP2 x LSL-Gq-DREADD control and ELA males (n = 8-10 mice/group; F_1,16_ = 6.780, p = 0.019, main effect of rearing x treatment, followed by Fisher’s LSD *post-hoc* tests: vehicle CTL vs vehicle ELA males p = 0.041; vehicle ELA vs. CNO ELA males p = 0.027; 2-way RM ANOVA).

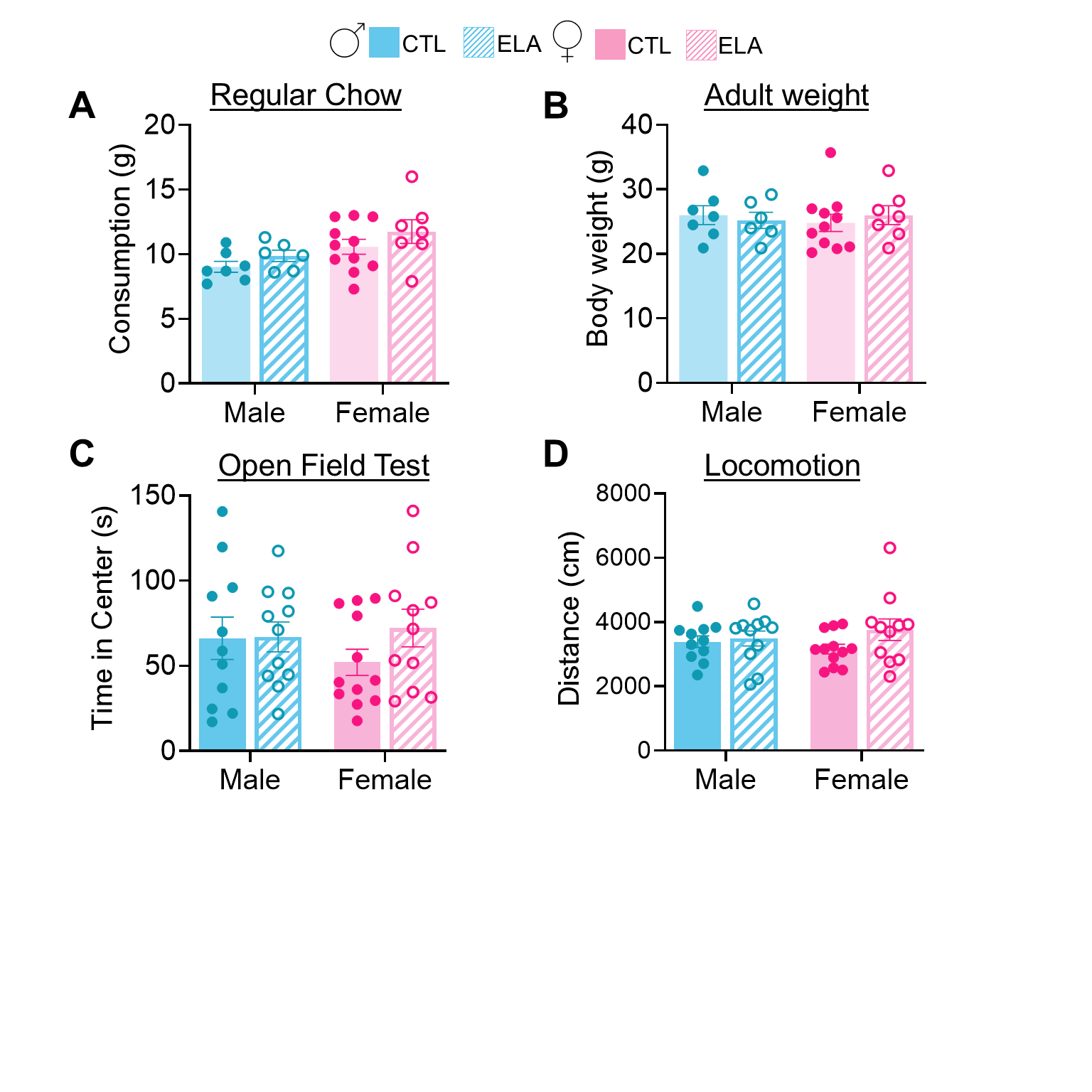

**Extended Data Figure 3. ELA does not influence anxiety, locomotion, regular chow consumption or adult body weight.** (**A**) Standard chow consumption across 72 hours does not differ between CTL and ELA males or CTL and ELA females (n = 6-11 mice/group; F_1,27_ = 0.060, p = 0.809, main effect of sex x rearing; F_1,27_ = 6.808, p = 0.015, main effect of sex; F_1,27_ = 2.389, p = 0.134, followed by Holm-Šídák's multiple comparisons test for Sex: male control vs female control, p = 0.134; male ELA vs female ELA, p = 0.134; 2-way ANOVA). (**B**) Adult body weight is not significantly different between control and ELA males or females (n = 6-11 mice/group; F_1,27_ = 0.484, p = 0.492, main effect of sex x rearing; F_1,27_ = 0.016, p = 0.899, main effect of sex; F_1,27_ = 0.016, p = 0.889; 2-way ANOVA). (**C**) Time in the center of the Open Field does not differ between groups (n =11-12 mice/group; F_1,41_ = 0.919, p = 0.343, main effect of sex x rearing; F_1,41_ = 0.195, p = 0.661, main effect of sex; F_1,41_ = 1.07, p = 0.306, main effect of rearing; 2-way ANOVA). (**D**) Distance traveled in the Open Field does not differ between groups (n =11-12 mice/group; F_1,41_ = 1.193, p = 0.281, main effect of sex x rearing; F_1,41_ = 0.008, p = 0.930, main effect of sex; F_1,41_ = 2.320, p = 0.135, main effect of rearing; 2-way ANOVA).

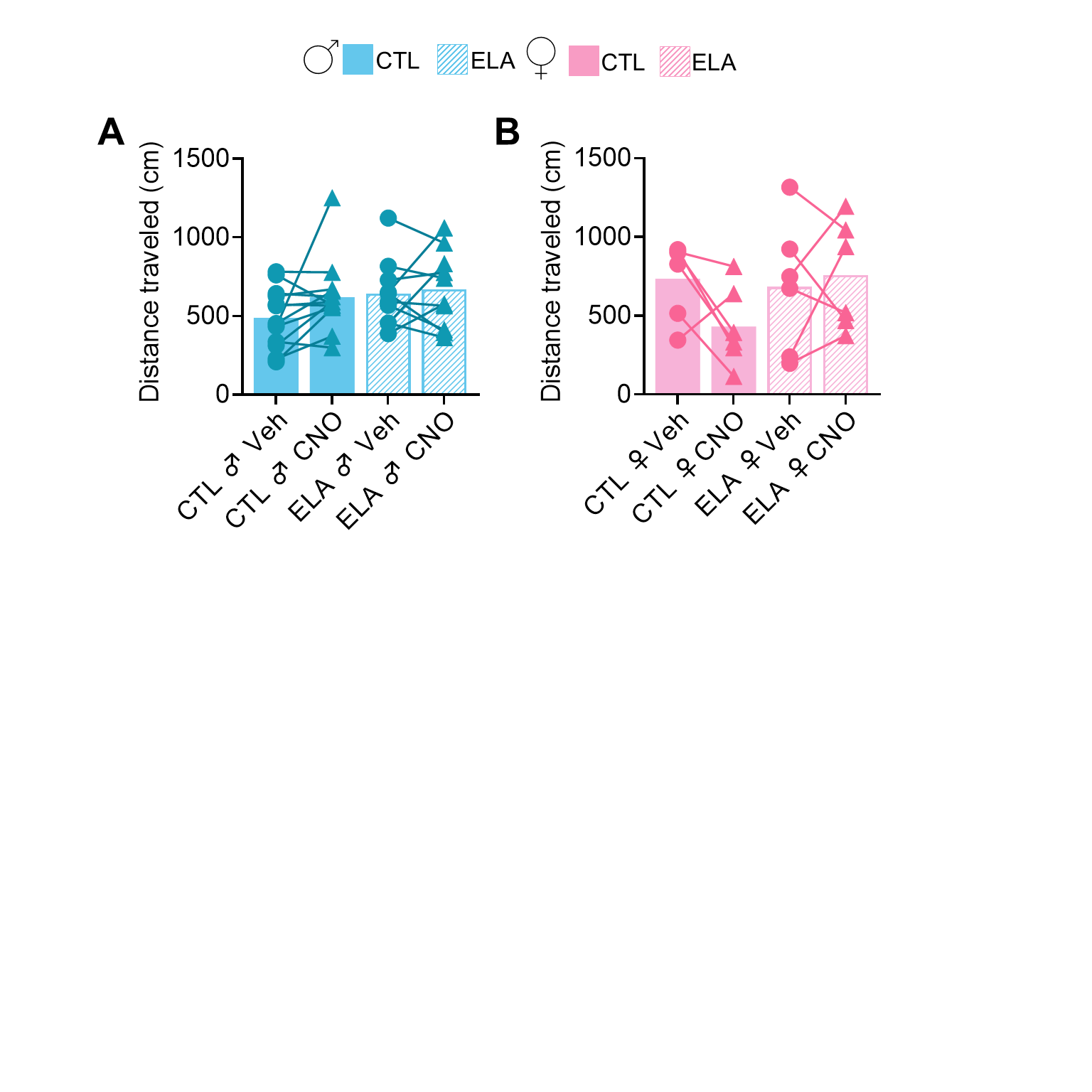

**Extended Data Figure 4. Intra-PVT CNO infusion and rearing do not influence locomotion.** (**A**) Neither ELA nor intra-PVT CNO microinfusion influence locomotion over the course of 3 minutes in males (n = 10-13 mice/group; F_1,21_ = 1.072, p = 0.312, main effect of rearing x treatment; F_1,21_ = 1.744, p = 0.201, main effect of rearing; F_1,21_ = 2.437, p = 0.133, main effect of treatment; 2-way RM ANOVA) (**B**) or females (n = 6 mice/group; ; F_1,10_ = 2.689, p = 0.132, main effect of rearing x treatment; F_1,10_ = 0.850, p = 0.378, main effect of rearing; F_1,10_ = 1.003, p = 0.340, main effect of treatment; 2-way RM ANOVA). While the 3-minute time frame analyzed for locomotion is brief, it coincides with the relevant behavior examined (i.e., sex cue interaction). A longer test would be required for general effects on locomotion.

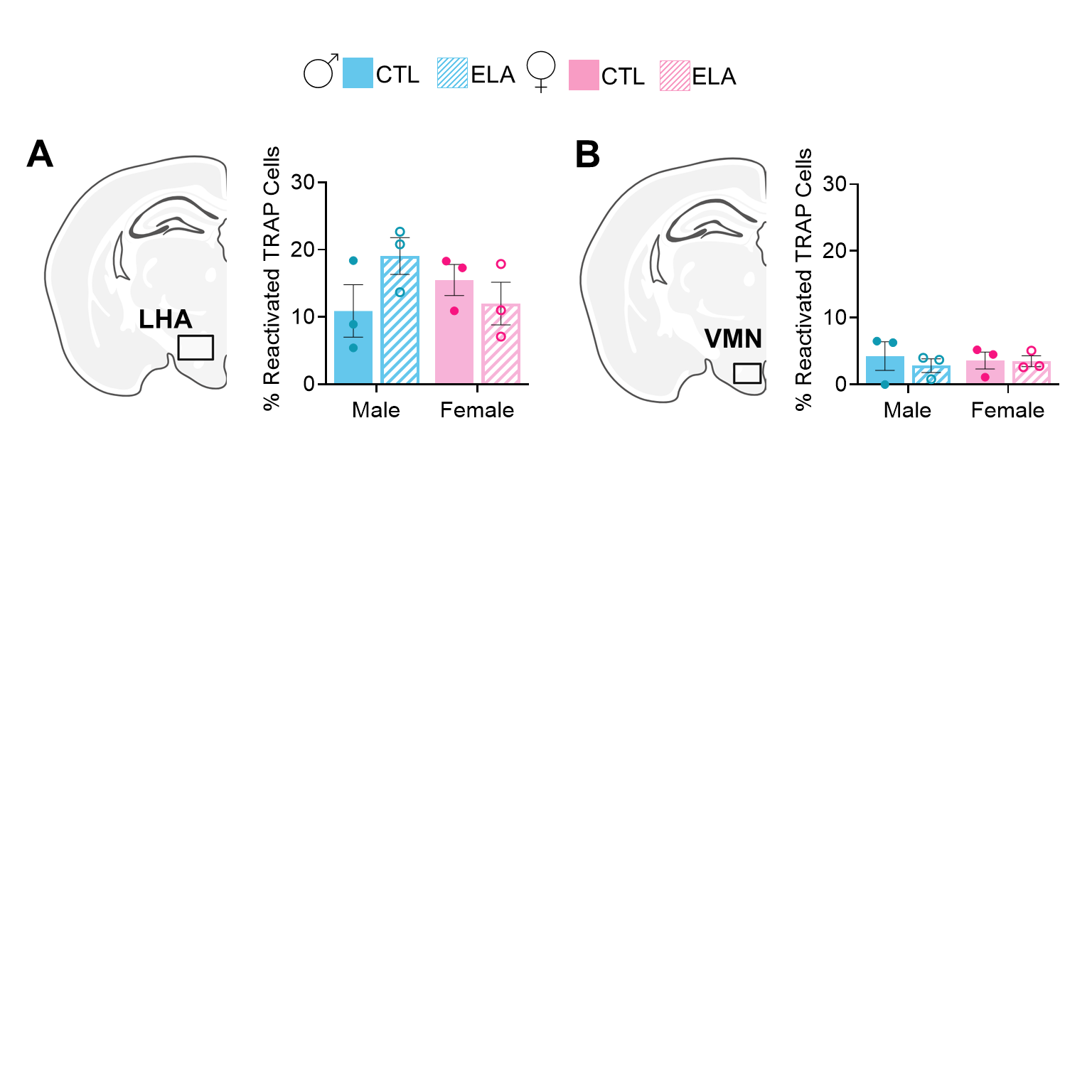

**Extended Data Figure 5. TRAP reactivation does not differ between control and ELA mice in the lateral hypothalamus or ventromedial hypothalamus.** (**A**) Neither sex nor rearing influenced the percent of reactivated TRAP neurons in the LHA (n = 3 mice/group; F_1,8_ = 3.589, p = 0.095, main effect of sex x rearing; F_1,8_ = 0.160, p=0.699, main effect of sex; F_1,8_ = 0.574, p = 0.470, main effect of rearing; 2-way ANOVA) (**B**) or VMH (n = 3 mice/group; F_1,8_ = 0.227 , p = 0.647, main effect of sex x rearing; F_1,8_ = 0.00, p>0.999, main effect of sex; F_1,8_ = 0.300, p = 0.599, main effect of rearing; 2-way ANOVA). Abbreviations: LHA, lateral hypothalamic area; VMN, ventromedial hypothalamic nucleus

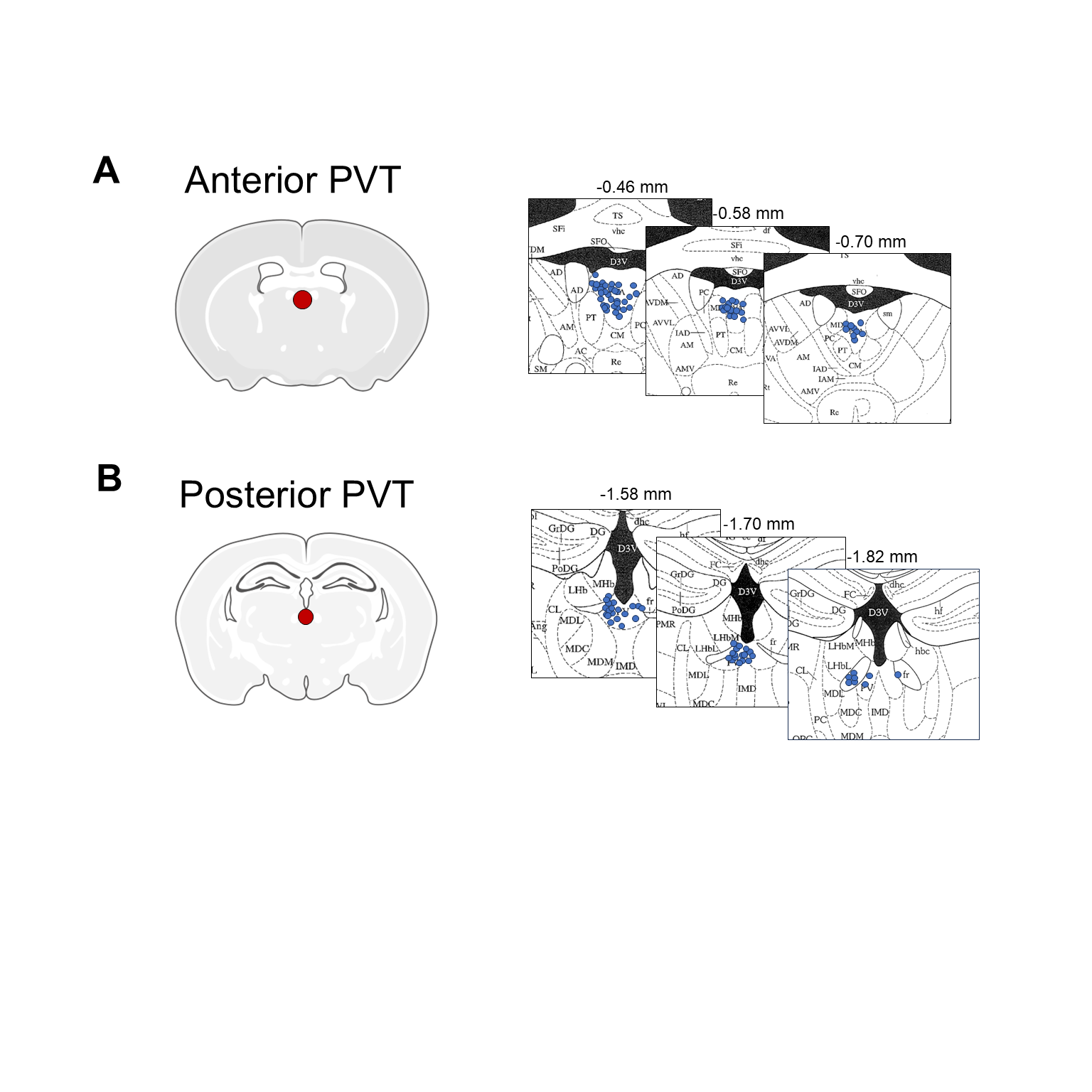

**Extended Data Figure 6. Representative map of PVT injection locations for DREADD experiments** (**A**) Mapped representative terminal points of intra-aPVT treatment infusion cannulas at three points along the rostrocaudal axis (shown as mm from Bregma). (**B**) Terminal points of intra-pPVT treatment infusion cannulas at three points along the rostrocaudal axis (shown as mm from Bregma).
